# Microbial community composition is affected by press, but not pulse, seawater intrusion

**DOI:** 10.1101/2020.04.07.030791

**Authors:** Courtney Mobilian, Nathan I. Wisnoski, Jay T. Lennon, Merryl Alber, Sarah Widney, Christopher B. Craft

**Affiliations:** School of Public and Environmental Affairs, Indiana University, Bloomington, Indiana, USA; Department of Biology, Indiana University, Bloomington, Indiana, USA; Department of Marine Sciences, University of Georgia, Athens, GA, USA

**Keywords:** Microbial diversity, community composition, seawater intrusion, disturbance, pulse, press, tidal freshwater marsh, climate change

## Abstract

Tidal freshwater marshes (TFMs) are threatened by seawater intrusion, which can affect microbial communities and alter biogeochemical processes. Here, we report on Seawater Addition Long Term Experiment (SALTEx), a manipulative field experiment that investigated continuous (press) and episodic (pulse, 2 months/yr) inputs of brackish water on microbial communities in a TFM. After 2.5 years, microbial diversity was lower in press treatments than in control (untreated) plots. Sulfate reducers increased in response to both press and pulse treatments whereas methanogens did not differ among treatments. Our results suggest that microbial communities in TFMs are resilient to episodic events, but that continuous seawater intrusion may alter bacterial diversity in ways that affect ecosystem functioning.

**Scientific Significance Statement:** Sea level rise and seawater intrusion threaten tidal freshwater marshes (TFMs) and the important ecosystem services they provide. Intrusion of seawater in TFMs can occur across a range of timescales, such as episodic events, like storm surges or drought, or continuous intrusion as a result of rising sea level. The effects of these stressors on TFM microbial communities are not well understood. Our multi-year field manipulation of brackish water inputs revealed that microbial communities were resilient to short-term pulses of salinity whereas continuous seawater intrusion led to reduced microbial diversity along with changes in relative abundance of key functional groups. Such alterations may diminish the ability of TFMs to sequester carbon and cycle nutrients.

## INTRODUCTION

Tidal freshwater marshes (TFMs) provide important ecosystem services such as carbon (C) sequestration (Loomis and Craft 2010), water quality improvement through the removal of nutrients such as nitrogen (N) and phosphorus (P) (Gribsholt et al. 2005; Neubauer et al. 2005), and protection from storm surges (Barbier et al. 2013), while also supporting high biodiversity of plants, animals, and microbial communities (Odum 1988). However, TFMs are threatened by climate change and rising seas, which are predicted to increase an additional 0.4-1.2 meters by the end of the century (Horton et al. 2014). Sea level rise is expected to lead to seawater intrusion into TFMs and tidal freshwater forests, converting them to brackish marshes, mud flats, or open water (DeLaune et al. 1994; Herbert et al. 2015). TFMs will also experience episodic (pulse) seawater intrusion owing to more frequent and longer periods of drought, increased occurrence of storm surges, and decreased freshwater inputs from rivers (van Vliet et al. 2013). It is unclear how these at-risk ecosystems will respond to increased salinity from such continuous and episodic incursions of seawater.

Salinity alters biogeochemical processes in tidal marshes, due in part to changes in both the activity and composition of microbial communities (Reed and Martiny 2013). Seawater intrusion modifies the composition and availability of electron acceptors, such as sulfate (Capone and Kiene 1988). Higher sulfate levels could lead to major shifts in the dominant microbial functional groups in TFMs, such as an increased abundance of sulfate reducers and a decrease in methanogens in response to altered redox conditions (Weston et al. 2006). While some microbial communities are thought to be resistant and resilient to environmental change (Shade et al. 2012), it is unclear how TFM microbial communities will respond to seawater inputs in the coming decades.

Microbial responses may depend on the amount, timing, and duration of seawater intrusions. For example, reciprocal transplants of salt marsh soils into a freshwater habitat revealed that, after 40 days, abundance of methanogens (*mcrA*) increased, but the abundance of sulfate reducers (*dsrA*) was unaffected (Morrissey and Franklin 2015). However, after 40 days, communities remained more phylogenetically related to their “home” environment compared to the “away” environment, suggesting that microorganisms are resilient to short-term perturbations (Morrissey and Franklin 2015). In a one-year study, transplanting TFM soils into a mesohaline environment led to an increased abundance of sulfate reducing bacteria (Dang et al. 2019). Together, these studies demonstrate that microorganisms can be sensitive to changes in salinity, but that the responses may depend on the duration of the perturbation.

In this study, we evaluated changes in the microbial community in response to episodic (pulse) and continuous (press) seawater intrusions by conducting a field-scale manipulation. In our experiment, replicated TFM plots (n=6) received press or pulse additions of brackish water, freshwater, or were left untreated (control). We hypothesized that microbial diversity would differ between plots receiving continuous and episodic seawater intrusion. We also hypothesized there would be increased abundance of sulfate reducers and decreased abundance of methanogens in plots receiving seawater additions relative to plots not receiving seawater additions.

## METHODS

### Site Description and Experimental Design

The experiment, Seawater Addition Long-Term Experiment (SALTEx), was conducted in a TFM on the Altamaha River Georgia, USA, as part of the Georgia Coastal Ecosystems Long Term Ecological Research (GCE-LTER) project (http://gce-lter.marsci.uga.edu/) (Figure S1). The site is dominated by giant cutgrass, *Zizaniopsis miliacea* Michx, and experiences twice-daily tidal inundations of freshwater with an average flooding depth of 25 cm at high tide (Widney et al. 2019). Details of the long-term experiment can be found elsewhere (Herbert et al. 2018, Widney et al. 2019), but briefly, we established 2.5 × 2.5 m replicated (n = 6 per treatment) plots at the site, which were randomly assigned to one of the following treatment groups: Control, Fresh, Pulse salinity, and Press salinity. Beginning in April 2014, Press plots were dosed four times per week with approximately 265 L of treatment water (∼15 ppt salinity), consisting of an equal mixture of seawater and fresh river water. Pulse plots were dosed four times per week with treatment water in September and October, during natural low river flow and dosed with fresh river water the remaining 10 months of the year. Fresh plots were dosed with fresh river water four times per week. Control (untreated) plots did not receive brackish water but, like the other treatments, were regularly inundated by the tides.

### Sample Collection and Environmental Variables

After treatment additions for 2.5 years, we collected soil samples (0-10 cm) from each of the four replicate plots from each treatment group in October 2016. Soils were placed in a cooler and shipped frozen to Indiana University where they were stored in a -80°C freezer until ready for DNA extraction.

We also measured porewater ammonium (NH_4_^+^), nitrate (NO_3_^-^), dissolved reactive phosphorus (DRP), and soil surface temperature quarterly, including October 2016, to identify potential drivers of community structure in each treatment. Porewater results and details on methods are described in more detail elsewhere (Herbert et al. 2018; Widney et al. 2019).

### Microbial Characterization

We characterized bacterial and archaeal composition using 16S rRNA amplicon sequencing. After extracting DNA from each sample using a MoBio PowerSoil DNA extraction kit (Carlsbad, CA), we amplified the V4 region of the 16S rRNA gene using 5PRIME HotMasterMix and 515F and 806R primers with customized Illumina sequencing adapters and unique sample barcodes following conditions described in detail elsewhere (Daum 2017). Amplicons were then pooled at approximately equal molar concentrations after quantification using a Roche LightCycler 480 real-time PCR instrument. The pooled sample was then sequenced on the MiSeq sequencing platform with a v3 600 Reagent kit following a 2×300 indexed run recipe at the at the Joint Genome Institute in Walnut Creek, California. Raw sequences were processed using the iTagger v. 2.2 pipeline (https://bitbucket.org/berkeleylab/jgi_itagger/src/itagger2/) and USEARCH (v. 9.2). Briefly, paired end reads were merged and quality filtered using expected error filtering. The resulting sequences were then incrementally clustered into operational taxonomic units (OTUs) starting at 99% identity and sequentially increasing the clustering radius by 1%. Finally, OTUs were classified using the Ribosomal Database Project (RDP) reference. The final OTU table and metadata can be found in the Zenodo archive (Craft et al. 2020), and raw sequence data are available at the NCBI Sequence Read Archive (BioProject PRJNA611801).

### Diversity Analysis

To assess microbial responses to seawater intrusion, we compared patterns of α- and β-diversity among treatments. We performed rarefaction on the community data, subsampling communities to 143,105 reads, and then relativized operational taxonomic unit (OTU) abundances to the total number of reads per sample. We characterized within-sample (α) diversity as the effective number of OTUs by taking the exponential of Shannon’s index (i.e., Hill number with degree = 1), which improves comparisons among groups (Jost 2006). We used ANOVA to compare differences in α-diversity among treatments, followed by Tukey’s Honest Significant Difference method to generate confidence intervals for group differences. To explain differences in community structure among treatments (i.e., β-diversity), we first transformed OTU relative abundances with the Hellinger transformation, and then used permutational multivariate ANOVA (PERMANOVA) to determine the significance of the treatment effects. We used redundancy analysis (RDA) to quantify the importance of individual environmental variables (porewater salinity, sulfides, NH_4_^+^, NO_3_^-^, and DRP) for explaining community differences among treatments.

We also analyzed the responses of two key functional groups to the experimental manipulations. First, we classified potential sulfate reducers as a subset of 16S rRNA sequences belonging to the following orders in the δ-Proteobacteria: Desulfuromonadales, Desulfarculales, Desulfobacterales, Desulfovibrionales, Desulfurellales, while recognizing that this may be an incomplete estimate of sulfate reducers. Second, we classified potential methanogens based on the summed relative abundances of archaeal sequences belonging to the following orders: Methanobacteriales, Methanomicrobiales, Methanosarcinales, and Methanocellales. We used ANOVA and Tukey’s method to determine the significance and effects of experimental treatments on the relative abundances of these taxonomic groups in the communities. All analyses were completed using the R environment (v. 3.6.0) and the vegan R package (Oksanen et al. 2019).

## RESULTS

### Microbial Composition

Although Pulse additions had no effect on α-diversity relative to the Control treatment (Tukey HSD, p = 0.441), Press inputs of seawater reduced α-diversity by 25% (ANOVA, *F*_*3,11*_ = 4.98, p = 0.017, Figure 1). Seawater additions also affected microbial community composition (PERMANOVA, *F*_*3,11*_ = 4.43, p = 0.001, r^2^ = 0.55). Based on the ordination plot from our RDA, community composition in Press treatments diverged from the composition of other treatments, while composition in the Pulse treatments was more similar to the Control and Fresh treatments (Figure 2). Microbial composition in the Pulse treatment was associated with higher porewater salinity, sulfides, and inorganic nitrogen (NO_3_^-^) than control and freshwater treatments (Table 1, Figure 2). Porewater in the Pulse treatments experienced transient changes during dosing, such as increased salinity and sulfides, that returned to background levels after dosing ceased (Figure 3) (Widney et al. 2019). Our RDA also revealed that, while also having increased salinity, the Press treatment was distinguished from Pulse and other treatments by higher porewater NH_4_^+^, dissolved reactive phosphorus (DRP), and soil surface temperature (Table 1, Figure 2). Continuous (press) salinity also led to more persistent changes in porewater chemistry, including increased salinity, ammonium, and sulfides for the duration of the experiment (Figure 3) (Widney et al. 2019).

**Table 1.** Redundancy Analysis of microbial composition vs. environmental variables. Correlations between environmental vectors and ordination axes are presented in the RDA1 and RDA2 columns. R^2^ values and p-values were determined by permutation (n = 999) using the envfit function in the vegan R package.

| | RDA1 | RDA2 | $r^2$ | $p$ |
| --- | --- | --- | --- | --- |
| DRP | 0.96 | 0.27 | 0.54 | **0.008** |
| $\text{NH}_4^+$ | 0.93 | 0.37 | 0.80 | **0.001** |
| $\text{NO}_3^-$ | 0.66 | -0.75 | 0.10 | 0.537 |
| Sulfide | 0.63 | -0.78 | 0.33 | 0.062 |
| Salinity | 0.55 | -0.83 | 0.67 | **0.001** |
| Soil Surface Temperature | 0.99 | 0.16 | 0.87 | **0.001** |

**Figure 1.**
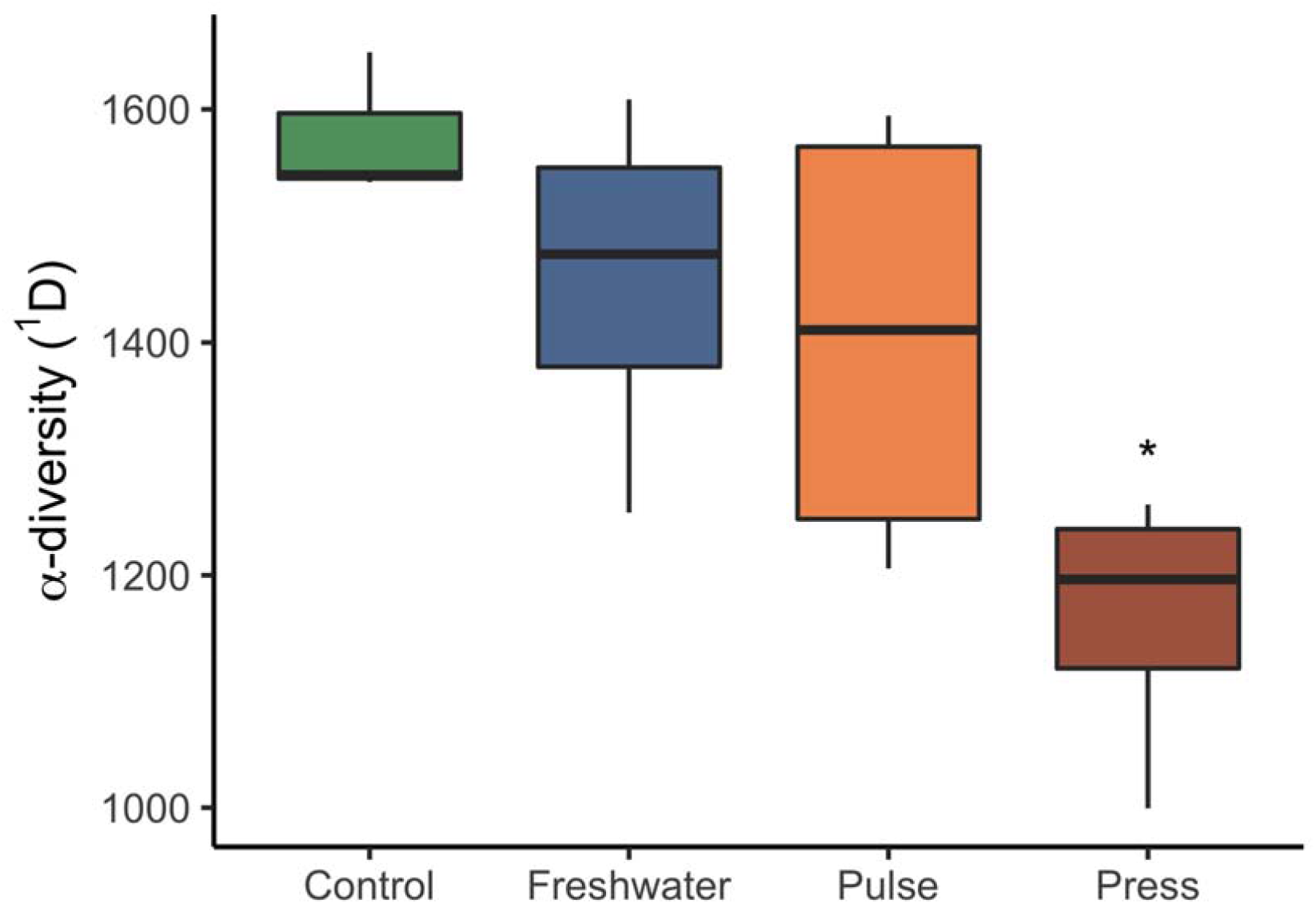
Alpha diversity of the total DNA community in each treatment group. * indicates Press is different from Control (p = 0.017).

**Figure 2.**
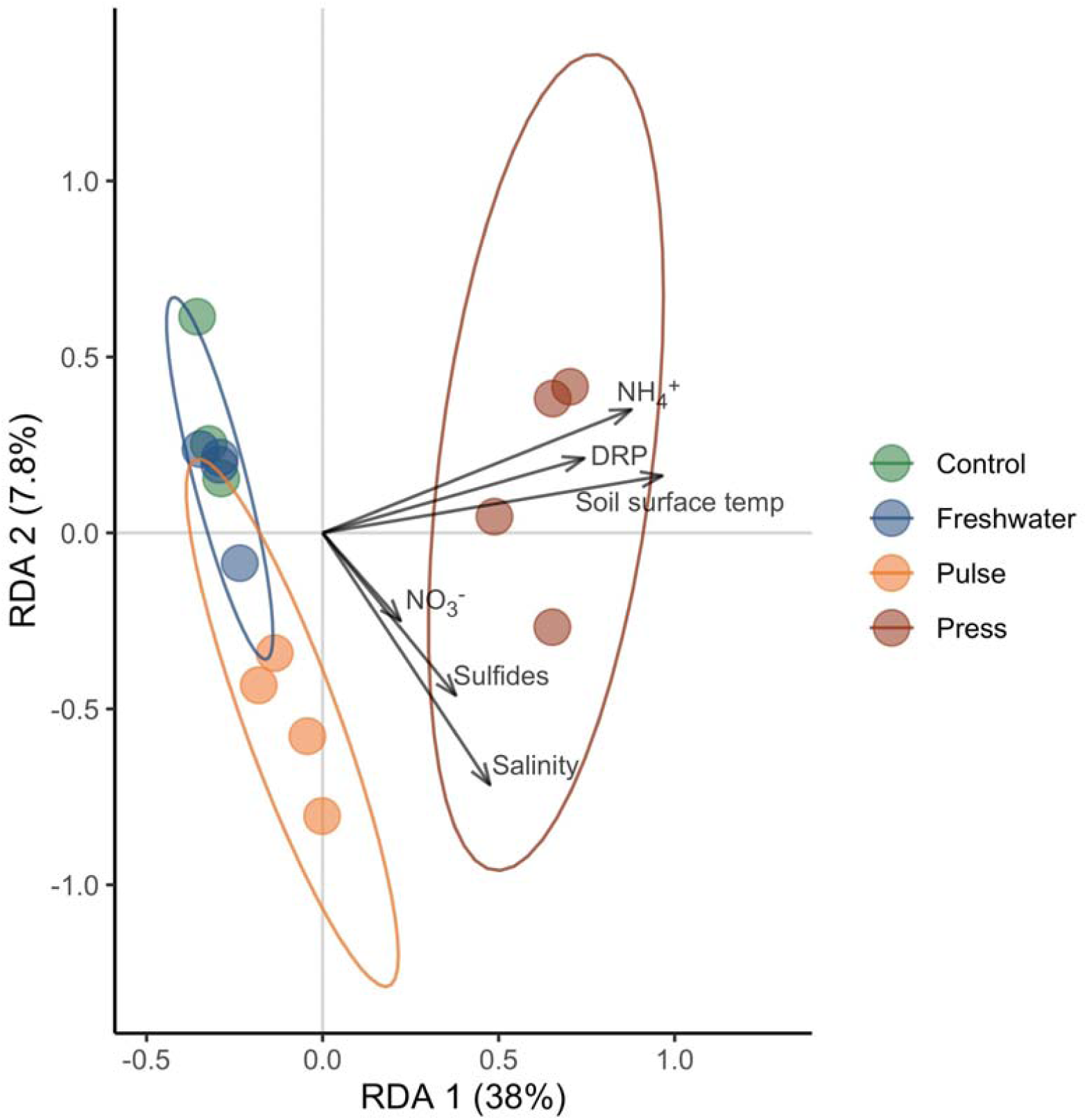
Redundancy analysis of microbial community structure with vectors depicting environmental variables along axes RDA1 and RDA2. Environmental data came from Herbert et al. 2018 and Widney et al. 2019.

**Figure 3.**
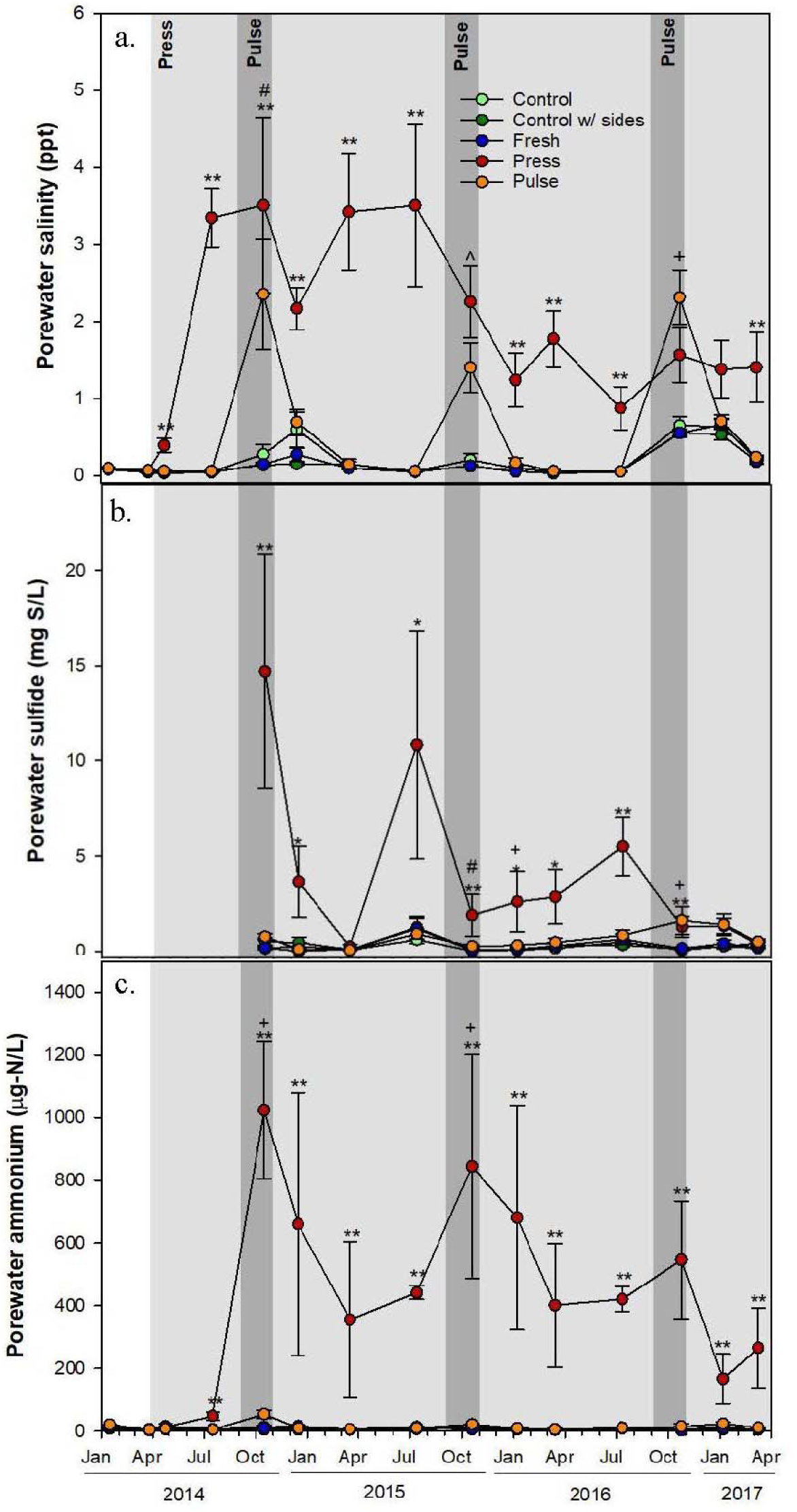
Porewater salinity (a), sulfide (b), and ammonium (b) concentrations (means ± SE) of treatments. Light gray shading indicates duration of Press treatment and darker gray shaded bars indicate the timing and duration of the Pulse treatment. Microbial samples were collected in October 2016. ** = Press > other treatments (p<0.05); * = Press is greater than other treatments (p<0.10); # = Press > other treatments except Pulse (p<0.05); + = Pulse > other treatments except Press (p<0.05); ^ = Press and Pulse > other treatments (p<0.05). Figure modified from Widney et al. 2019.

Seawater manipulations also had an effect on some microbial functional groups. For example, the relative abundance of sulfate reducers was nearly double in the Press plots (6.5%, Tukey HSD, p = 0.0004) than in Control treatments (3.5%) (ANOVA, *F*_*3,11*_ = 15.34, p = 0.003, Figure 4a). In addition, sulfate reducers were enriched in the Pulse treatment (6%) compared to the Control treatment (3.5%) (Tukey HSD, p = 0.0078) (Figure 4a). The relative abundance of sulfate reducers was also greater in the Press (Tukey, HSD, p = 0.0019) compared to the Fresh treatment and marginally greater in the Pulse compared to the Fresh treatment (Tukey HSD, p = 0.054) (Figure 4a). Relative abundance of sulfate reducers did not differ between Press and Pulse treatments (Tukey HSD, p = 0.23). Contrary to our prediction, the relative abundance of methanogens, which ranged from 0.5 to 1.5%, did not differ among treatments (ANOVA, *F*_*3,11*_ = 0.494, p = 0.694) (Figure 4b).

**Figure 4.**
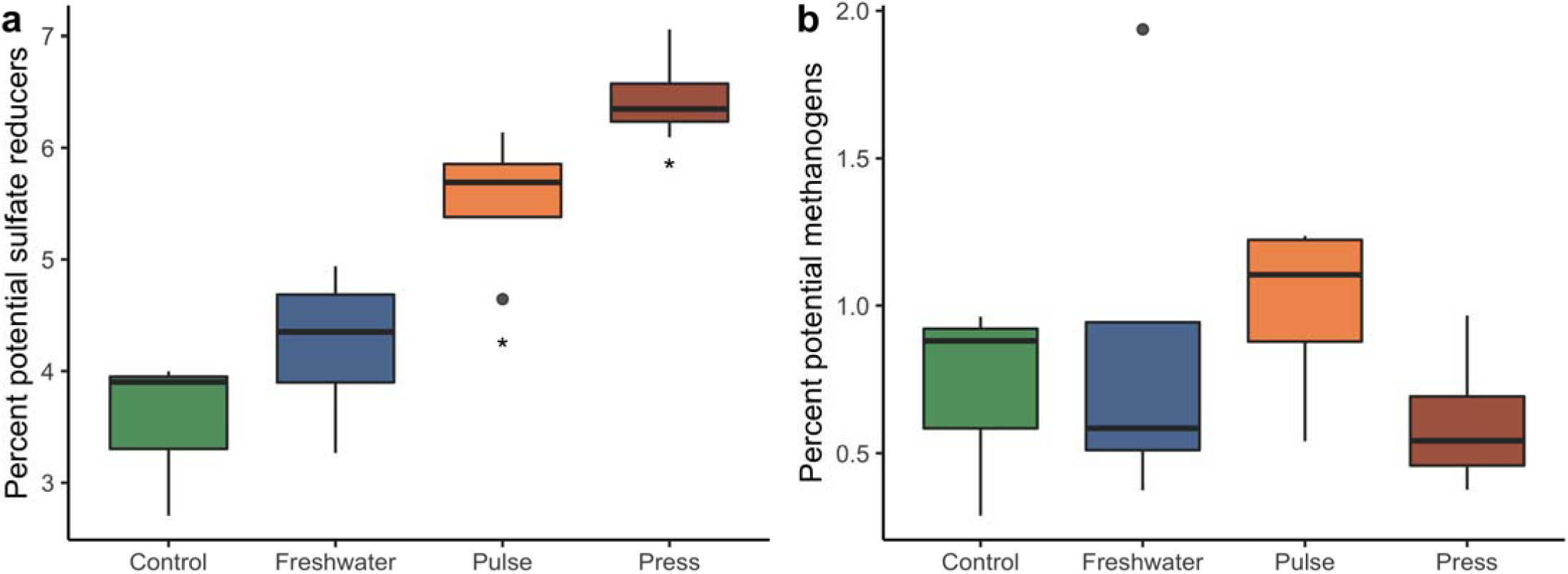
(a).Sulfate reducer abundance and (b). methanogen abundance based on the total DNA community. * indicates Press is different from Control (p = 0.0004) and Fresh (p = 0.0019) and Pulse is different from Control (p = 0.0078) and Fresh (p = 0.0539).

## DISCUSSION

Salinity is an important driver of microbial community composition (Lozupone and Knight 2007; Morrissey et al. 2014), which can in turn affect biogeochemical processes including carbon sequestration (Weston et al. 2011; Neubauer et al. 2013; Morrissey et al. 2014). In this study, we observed reduced microbial diversity, altered microbial composition, and an increase in the relative abundance of sulfate reducing bacteria (SRB) in response to continuous (press) seawater intrusion for 2.5 years. In contrast, our results suggest that microbial communities in TFMs are less affected by episodic (pulse) seawater intrusion. Taken together, our study indicates that modified salinization regimes in TFMs may have implications for sulfur cycling, in particular increased potential for sulfate reduction, that emerge over longer time scales of seawater intrusion.

Microbial community composition in the Pulse plots was different from both the Control and the Press treatments (Figure 2). Relative to Control treatments, Pulse plots had a greater abundance of sulfate reducers (Figure 4a) but no difference in diversity (Figure 1). Other short-term studies have also reported no change in diversity in response to increased salinity (e.g., Edmonds et al. 2009 after 3 weeks of dosing, and Dang et al. 2019 after 1 year), similar to the response we observed in our Pulse plots. In response to Press seawater additions, SRB became even more common in the community (Figure 4a), and we were able to detect overall declines in microbial diversity (Figure 1) and more extreme shifts in community composition (Figure 2). Thus, our study suggests that the frequency and duration of seawater intrusion affect the diversity and structure of TFM microbial communities.

Changes in the abundances of SRB could affect community diversity. Consistent with our hypothesis, we found greater abundance of SRB in Press plots (Figure 4a), which had higher concentrations of porewater sulfide compared to other treatments (Figure 3) (Widney et al. 2019). Our results are also consistent with other studies (Weston et al. 2006, 2011; Dang et al. 2019), including a transplant study of freshwater marsh soils into a brackish marsh that found a slight increase in abundance of sulfate reducers (*dsrA*) (Dang et al. 2019). A shift in the microbial community towards SRB has the potential to negatively affect other members of the community because sulfate reduction can generate hydrogen sulfide that is toxic to more sensitive members of the community. Thus, long-term, chronic inputs of seawater that maintain elevated concentrations of porewater sulfide could favor SRB and the production of hydrogen sulfide, which contribute to the reduction in overall microbial diversity.

Sulfate is an electron acceptor that can support sulfate reduction and this reaction is more thermodynamically favorable than methanogenesis. Therefore, we predicted that increases in SRB would be accompanied by a decrease in the relative abundance of methanogens. We did not observe this pattern in our data. As methanogenesis relies on decomposition of organic matter, other studies have indicated that methanogen abundances may be influenced by carbon availability (DOC and soil organic carbon) (Yuan et al. 2016; Dang et al. 2019). Although previous work from our experiment observed lower methane emissions in the Press treatment, there was little variation in methane production or DOC concentration across treatments during the month of October 2016 when the samples from the current study were collected (Herbert et al. 2018) suggesting that C limitation constrained the abundances of methanogens across all treatments. Sulfate-mediated anaerobic oxidation of methane (AOM), where sulfate is used as an electron acceptor for AOM, has been shown to be important in tidal freshwater sediments of the Altamaha River (Segarra et al. 2015) and could account for reduced CH_4_ emissions from our Press plots.

Microbial community responses to seawater intrusion may also depend on the concentrations of other nutrients. For example, in a lab microcosm experiment, where wetland sediments were exposed to both increasing salinity and nutrients (N, P), salinity alone increased bacterial diversity, whereas N additions (nitrate, ammonium) and the combination of increased salinity, nitrogen and phosphorus decreased diversity (Jackson and Vallaire 2009). In our long-term field experiment, Press additions of brackish water resulted in not only elevated salinity but also higher porewater inorganic nitrogen (ammonium, nitrate) concentrations compared to the other treatments (Figure 3) (Herbert et al. 2018; Widney et al. 2019) which could explain the reduced α-diversity in the Press plots. α-diversity did not differ between Pulse and Control treatments and, while the Pulse plots experienced transient increases in salinity, porewater N did not increase as it did in the Press plots (Herbert et al. 2018; Widney et al. 2019).

Shifts in microbial community structure may also be facilitated by the immigration of taxa that accompanied the intrusion water. That is, seawater additions not only cause the mixing of freshwater and brackish environments, they also lead to the mixing of brackish and freshwater microorganisms, a phenomenon referred to as “community coalescence” (Rillig et al. 2015). The resulting community composition may depend on many factors, including (1) differences in environmental conditions, where seawater was added to a freshwater environment and (2) the temporal dynamics of mixing (Rillig et al. 2015). For example, Pulse seawater additions temporarily modified the environment (for ∼2 months, Figure 3), but this treatment may have been insufficient to overcome the larger and more frequent additions of fresh river water and freshwater microorganisms during the natural tidal cycle. As a result, microbial communities in Pulse plots were only slightly different from the Control and Fresh treatments (Figure 2).

In conclusion, continuous seawater intrusion can reduce microbial diversity and alter community composition, which may be due to increases in porewater sulfate that favor an increase in abundance of sulfate reducers. Episodic seawater additions did not affect α-diversity but increased sulfate reducer abundance, suggesting that microbial communities of the Pulse treatment retain environmental and microbial characteristics of freshwater tidal marshes. The increase in sulfate reducers (and sulfate reduction) we observed in the Press plots may have implications for C and nutrient (P) cycles, such as increased C mineralization and phosphorus release. Furthermore, the decline in microbial diversity in Press plots may reduce the functional redundancy of these and other microbial processes.

## Supporting information

Metadata

## Data Availability Statement

Sequence data is available on the NCBI SRA (BioProject PRJNA611801). Data, metadata, and code used to generate results will be made publicly available on a Zenodo archive (Accession # and doi pending) of the GitHub repository https://github.com/LennonLab/MicroMarsh.

## Acknowledgements

We thank Ellen Herbert for collecting the soil samples for analysis and thank Steve Reynolds for extracting DNA.

## Supplemental Information

**Figure S1.**
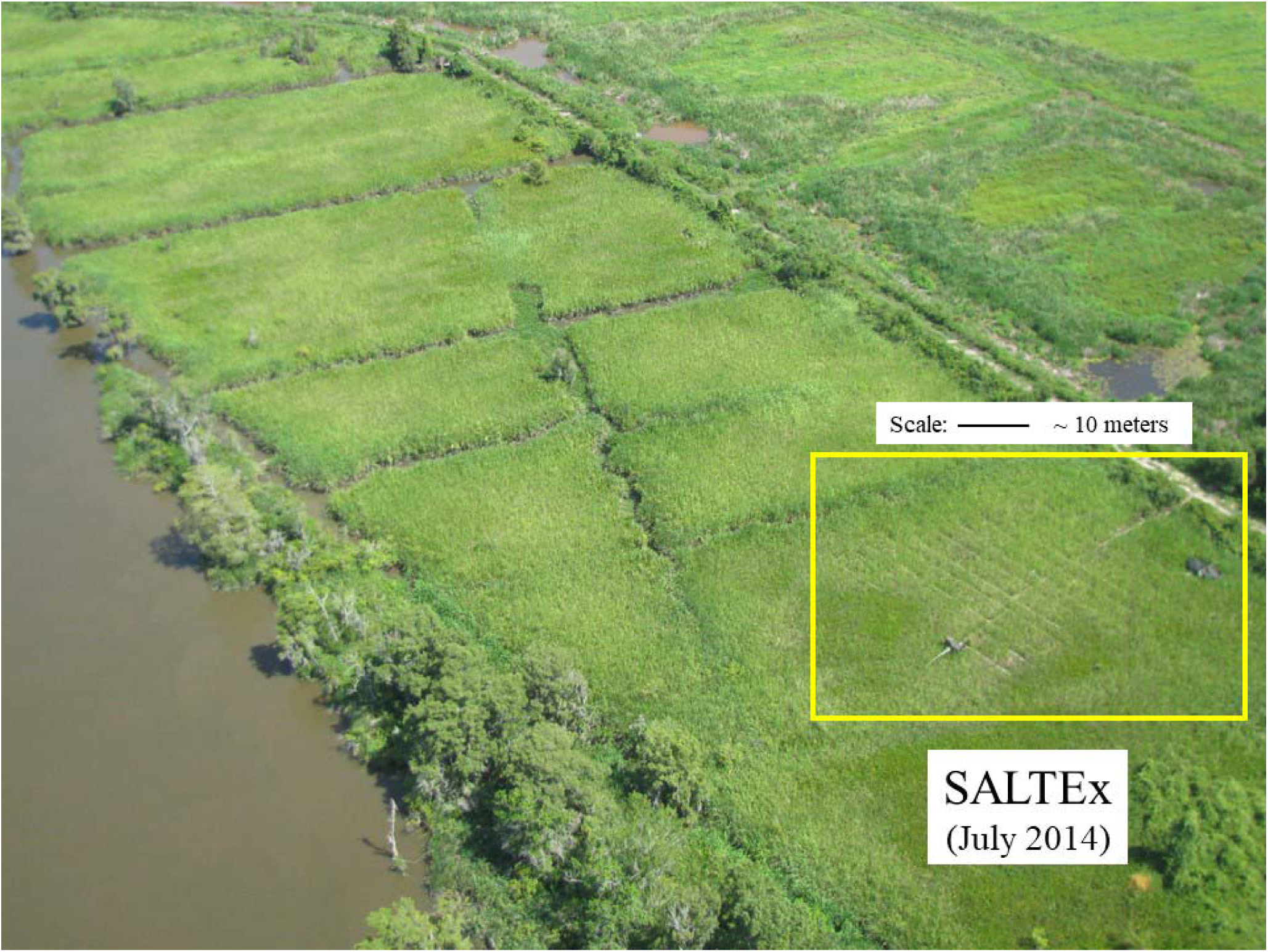
Aerial view of the SALTEx experimental site in July 2014.

