## Supplementary material for "Microbial community composition is affected by press, but not pulse, seawater intrusion": Metadata

### Metadata template for datasets of *L&O-Letters* articles

**Table 1.** Description of the fields needed to describe the creation of your dataset.

| | |
|---|---|
| **Title of dataset** | Soil microbial communities in a tidal freshwater marsh being experimentally dosed with episodic (pulse) and continuous (press) additions of seawater. |
| **URL of dataset** | Will be provided upon acceptance |
| **Abstract** | A large-scale saltwater intrusion field experiment was conducted in a tidal freshwater marsh to investigate the effects of continuous and episodic brackish water intrusion on soil microbial communities. The experimental field site, SALTEx (Seawater Addition Long-Term Experiment) is part of the Georgia Coastal Ecosystems (GCE) LTER and is located on the Altamaha River, GA. 2.5 m by 2.5 m replicated (n=6 per treatment) plots were exposed to continuously high salinity (press plots) or episodic (pulse plots) high salinity (2 months/yr) or were untreated (control, fresh river water) starting in April 2014. Soil samples were collected in October 2016 and DNA was extracted to assess microbial community composition |
| **Keywords** | Sequence data, microbial community composition, seawater intrusion, pulse disturbance, press disturbance, tidal freshwater marsh, Georgia Coastal Ecosystems Long Term Ecological Research Project |
| **Lead author for the dataset** | Christopher Craft |
| **Title and position of lead author** | Principle Investigator |
| **Organization and address of lead author** | Indiana University, O'Neill School of Public and Environmental Affairs<br><br>Multidisciplinary Science Building II<br><br>Room 408<br><br>702 N. Walnut Grove St.<br><br>Bloomington, IN 47405 |
| **Email address of lead author** | |
| **Additional authors or contributors to the dataset** | Nathan Wisnoski, Jay Lennon, Courtney Mobilian |
| **Organization associated with the data** | Indiana University, Georgia Coastal Ecosystems Long Term Ecological Research (GCE LTER) project, and the Department of Energy (DOE) Joint Genome Institute |

| | |
|---|---|
| **Funding** | Principle Investigators: Christopher Craft and Jay Lennon<br><br>Funding:<br><br>U.S. Department of Energy Joint Genome Institute (JGI): Project ID 1178283.The work conducted by the U.S. Department of Energy Joint Genome Institute, a DOE Office of Science User Facility, is supported by the Office of Science of the U.S. Department of Energy under Contract No. DE-AC02-05CH11231.<br><br>National Science Foundation's Long-Term Ecological Research (LTER) program (Georgia Coastal Ecosystems LTER, OCE 9982133).<br><br>National Science Foundation (1442246, J.T.L.) and a US Army Research Office Grant (W911NF-14-1-0411, J.T.L.) |
| **License** | CCBY |
| **Geographic location – verbal description** | The study site, Seawater Addition Long-Term Experiment (SALTEx), is a tidal freshwater marsh on the Altamaha River near Darien, Georgia, USA |
| **Geographic coverage bounding coordinates** | 31°20′16″ N, 81°27′52″ W |
| **Time frame - Begin date** | April 2014 |
| **Time frame - End date** | October 2016 |
| **General study design** | The study site, Seawater Addition Long-Term Experiment (SALTEx), a TFM on the Altamaha River Georgia, USA, is part of the Georgia Coastal Ecosystems Long Term Ecological Research (GCE-LTER) project (http://gce-lter.marsci.uga.edu/). The site is dominated by giant cutgrass, Zizaniopsis miliacea Michx, and experiences twice daily tidal inundations of freshwater with an average flooding depth of 25 cm at high tide. 2.5 x 2.5 meter plots were established at the site and randomly assigned to a treatment group: Control, Fresh, Pulse salinity, and Press salinity, each replicated six times. Beginning in April 2014, Press plots were dosed 4 times per week with approximately 265 liters of treatment water (~15 ppt salinity), consisting of an equal mixture of seawater and fresh river water. Fresh plots were dosed with fresh river water 4 times per week. Pulse plots were dosed 4 times per week with treatment water in September and October, during natural low river flow then dosed with fresh river water the remaining 10 months of the year. |

| | |
|---|---|
| **Methods description** | Experimental plots were exposed to continuously high salinity (press) or episodic (pulse) high salinity (2 months/yr) for 2.5 years. Press plots were dosed 4 times per week with approximately 265 liters of treatment water (~15 ppt salinity), consisting of an equal mixture of seawater and fresh river water. Fresh plots were dosed with fresh river water 4 times per week. Pulse plots were dosed 4 times per week with treatment water in September and October, during natural low river flow then dosed with fresh river water the remaining 10 months of the year. Soil samples (0-10 cm depth) were collected in October 2016 and DNA was extracted for analysis of microbial communities. We also measured porewater ammonium ($NH_4^+$), nitrate ($NO_3^-$), dissolved reactive phosphorus (DRP), sulfide, sulfate, salinity, chloride, dissolved organic carbon (DOC) and soil surface temperature quarterly, including October 2016, to identify potential drivers of community structure in each treatment. |
| **Field methods** | **Microbial data:** We collected soil samples (0-10 cm) from each of the four replicate plots from each treatment group in October 2016, 2.5 years after dosing was initiated. Soils were placed in a cooler and shipped frozen to Indiana University where they were then stored at -80°C until ready for DNA extraction.<br><br>**Environmental data:** Porewater was collected from two wells (10-35 cm below the soil surface) within each plot two tidal cycles after treatment water was added. Porewater was collected in a 500 mL acid-washed Nalgene bottle to get a composite sample. For sulfide samples, 20 mL was taken from the composite sample and combined with 20 mL of Orion sulfide antioxidant buffer (SAOB) in a 50 mL centrifuge tube.<br><br>The composite sample was then divided to 125 mL acid-washed Nalgene bottles for analyses of nutrients and ion chromatography. After collection samples were immediately placed on ice and frozen until analysis.<br><br>Soil surface temperatures were collected monthly using an infrared thermometer. Four measurements were collected from each plot and then averaged for each plot. Temperature measurements were taken when there was little or no standing water in the plots. |

| | |
|---|---|
| **Laboratory methods** | **Microbial data:** We characterized bacterial and archaeal composition using 16S rRNA amplicon sequencing. After extracting DNA from each sample using a MoBio PowerSoil DNA extraction kit (Carlsbad, CA), we amplified the V4 region of the 16S rRNA gene using 5PRIME HotMasterMix and 515F and 806R primers with customized Illumina sequencing adapters and unique sample barcodes following conditions described in detail elsewhere. Amplicons were then pooled at approximately equal molar concentrations after quantification using a Roche LightCycler 480 real-time PCR instrument. The pooled sample was then sequenced on the MiSeq sequencing platform with a v3 600 Reagent kit following a 2x300 indexed run recipe at the at the Joint Genome Institute in Walnut Creek, California (doi).<br><br>**Environmental data:** Porewater nutrients were analyzed using the Lachat QuikChem 8500 Flow Injection Analysis system. Ammonium was analyzed using the QuikChem method 10-107-06-1-B (indophenol blue complex, MDL = 7 µg/L); Nitrate was analyzed using the QuikChem method 10-107-04-1-J (cadmium reduction/ EDTA red complex, MDL = 3 µg/L); and Phosphorus was analyzed using the QuikChem method 10-115-01-1-B (molybdate blue complex, MDL = 1 µg/L). Sulfate and chloride were analyzed with a Dionex ICS-2000 Ion Chromatograph with an AS11-HC analytical column (MDL = 0.5 mg/L for chloride and sulfate). Salinity was then converted from chloride data. DOC was analyzed using a Shimadzu TOC-VCPN analyzer (MDL = 150 µg C/L). Porewater sulfide was analyzed with an Orion Model 9616 Sure-Flow Combination Silver/ Sulfide Electrode with Optimum Results B filling solution. |
| **Analytical methods** | **Microbial data:** Raw sequences were processed using the iTagger v. 2.2 pipeline (https://bitbucket.org/berkeleylab/jgi_itagger/src/itagger2/) and USEARCH (v. 9.2). Briefly, paired end reads were merged and quality filtered using expected error filtering. The resulting sequences were then incrementally clustered into operational taxonomic units (OTUs) starting at 99% identity and sequentially increasing the clustering radius by 1%. Finally, OTUs were classified using the Ribosomal Database Project (RDP) reference. Included here is the OTU table used in this analysis. |
| **Taxonomic species or groups** | Bacteria and archaea |
| **Quality control** | Quality control for microbial data is detailed above in Analytical Methods.<br><br>For porewater data, blanks and standards were run every 10 samples to ensure accuracy and correct for instrument drift. |
| **Additional information** | |

**Table 2.** Data dictionary: description of the variables (i.e., columns) in EACH dataset.

Dataset filename: otu_table.csv
Dataset description: OTU reads for each plot

| **Column name** | **Description** | **Units** | **Code explanation** | **Data format** | **Missing data code** |
|---|---|---|---|---|---|
| *The name of the variable in the dataset; avoid special characters, dashes and spaces* | *A detailed description of the variable* | *Units the variable is measured in* | *If you use codes in your column, please explain each code, such as: LR = Little Rock Lake; A=sample; etc.* | *State exactly how the data are stored; for dates, state how it is formatted, including time zone, etc.* | *If data are missing, indicate how they are stored, such as NULL, NA, blank cell, etc.* |
| sample_ID | The ID linking each of the three tables included here. This is a code for the experimental treatments. | N/A | | | |
| otuX | Number of reads of OtuX in the data set. | Sequence reads following processing described above | OtuX is an arbitrary name. | Read numbers | |

Dataset filename: env.csv
Dataset description: Porewater and soil temperature data associated with each plot

| **Column name** | **Description** | **Units** | **Code explanation** | **Data format** | **Missing data code** |
|---|---|---|---|---|---|
| *The name of the variable in the dataset; avoid special characters, dashes and spaces* | *A detailed description of the variable* | *Units the variable is measured in* | *If you use codes in your column, please explain each code, such as: LR = Little Rock Lake; A=sample; etc.* | *State exactly how the data are stored; for dates, state how it is formatted, including time zone, etc.* | *If data are missing, indicate how they are stored, such as NULL, NA, blank cell, etc.* |
| sample_ID | The ID linking each of the three tables included here. This is a code for the experimental treatments. | N/A | | | |
| date | Date on which the sample was taken | mm/dd/yy | | mm/dd/yy; EST time zone | |

| | | | | | |
|---|---|---|---|---|---|
| treatment | Experimental treatment | factor | C = control<br>F = Freshwater<br>PR = Press<br>PU = Pulse | | |
| replicate | Experimental replicate | factor | reps 1- 6 | | some reps missing from data set |
| molecule | Is the read from an RNA or DNA sample | factor | RNA = active;<br>DNA = total | | |
| DRP | Concentration of dissolved reactive phosphorus in porewater | µg-P/L | | | |
| NH4 | Concentration of ammonium ($NH_4^+$) in porewater | µg-N/L | | | |
| NO2_3 | Concentration of nitrate ($NO_3^-$) in porewater | µg-N/L | | | |
| DOC | Concentration of dissolved organic carbon in porewater | µg-C/L | | | |
| Sulfides | Concentration of sulfides in porewater | µg-S/L | | | |
| Cl | Concentration of chloride ($Cl^-$) in porewater | mg/L | | | |
| SO_4_2 | Concentration of sulfate ($SO_4^{2-}$) in porewater | mg/L | | | |
| Salinity | Salinity of porewater | ppt | | | |
| Soil_surface_temp | Average soil surface temperature | °C | | | |

Dataset filename: tax.csv
Dataset description:  Taxonomic description of each OTU

| **Column name** | **Description** | **Units** | **Code explanation** | **Data format** | **Missing data code** |
|---|---|---|---|---|---|

| | | | | | |
|---|---|---|---|---|---|
| *The name of the variable in the dataset; avoid special characters, dashes and spaces* | *A detailed description of the variable* | *Units the variable is measured in* | *If you use codes in your column, please explain each code, such as: LR = Little Rock Lake; A=sample; etc.* | *State exactly how the data are stored; for dates, state how it is formatted, including time zone, etc.* | *If data are missing, indicate how they are stored, such as NULL, NA, blank cell, etc.* |
| OTU | OTU number corresponding to the otu_table.csv | N/A | | | |
| Taxonomy | Taxonomic description associated with each OTU | N/A | d: = domain; p: = phylum; c: = class; o: = order; f: = family; g: = genus | | |

Dataset filename: design.csv
Dataset description: This table serves as a key that links the experimental treatments for each sample with the OTU table and environmental table.

| **Column name** | **Description** | **Units** | **Code explanation** | **Data format** | **Missing data code** |
|---|---|---|---|---|---|
| *The name of the variable in the dataset; avoid special characters, dashes and spaces* | *A detailed description of the variable* | *Units the variable is measured in* | *If you use codes in your column, please explain each code, such as: LR = Little Rock Lake; A=sample; etc.* | *State exactly how the data are stored; for dates, state how it is formatted, including time zone, etc.* | *If data are missing, indicate how they are stored, such as NULL, NA, blank cell, etc.* |
| sampleID | The ID linking each of the three tables included here. This is a code for the experimental treatments. | NA | | | |
| date | Date of sample collection | NA | YYYY-MM-DD | YYYY-MM-DD, EST | |
| treatment | experimental treatment for each plot | NA | C = control<br>F = freshwater<br>PR = press<br>PU = pulse | | |
| replicate | which experimental replicate this sample is | NA | replicates could be in the range of 1-6 | | |
| molecule | Is this sample an RNA or DNA sample? | NA | DNA = “total community”<br>RNA = “active community” | | |

#### Notes and Comments:
